# A deficiency screen identifies genomic regions critical for sperm head-tail connection

**DOI:** 10.1101/2024.08.20.608819

**Authors:** Brian J. Galletta, Parthena Konstantinidou, Astrid D. Haase, Nasser M. Rusan

## Abstract

A stable connection between the sperm head and tail is critical for fertility in species with flagellated sperm. The head-tail coupling apparatus (HTCA) serves as the critical link between the nucleus (head) and the axoneme (tail) via the centriole. To identify regions of the *Drosophila melanogaster* genome that contain genetic elements that influence HTCA formation, we undertook a two part screen using the *Drosophila* deficiency (Df) kit. For this screen, we utilized a sensitized genetic background that overexpresses the pericentriolar material regulatory protein Pericentrin-Like Protein (PLP). We had previously shown that PLP overexpression (PLP^OE^) disrupts the head-tail connection in some spermatids, but not to a degree sufficient to reduce fertility. In the first step of the screen we tested for Dfs that in combination with PLP^OE^ cause a reduction in fertility. We ultimately identified 11 regions of the genome that showed an enhanced fertility defect when combined with PLP overexpression. In the second step of the screen we tested these Dfs for their ability to enhance the HTCA defect caused by PLP^OE^, finding six. We then tested smaller Dfs to narrow the region of the genome that contained these enhancers. To further analyze the regions of the genome removed by these Dfs, we examined the expression patterns of the genes within these Dfs in publicly available datasets of RNAseq of *Drosophila* tissues and snRNAseq of *Drosophila* testes. In total, our analysis suggests that some of these Dfs may contain a single gene that might influence HTCA formation and / or fertility, while others appear to be regions of the genome especially rich in testis-expressed genes that might affect the HTCA because of complex, multi-gene interactions.

**Article Summary:** We perform a genetic enhancer deficiency screen to uncover genomic regions required for proper sperm head-tail connection. We identified 6 regions and provide insight into these regions using publicly available RNA sequence data. Our data reveal that these regions are exceptionally rich in testes specific genes. Further analysis using small deficiencies resulted in two classes of enhancers: one class likely enhances head-tail connection by disrupting multiple genes, while the second class might house a single gene responsible for the reduction in fertility.

## Introduction

The flagellated sperm of many organisms can be divided into two major parts, the head and the tail. The head contains the nucleus and the genetic material to be passed on to the next generation. The tail contains the flagellum (or cilium), which provides sperm with the ability to swim. A central component of the linkage between the sperm head and tail is the centriole (in this context often also referred to as a basal body), which templates the flagellum from its distal end and attaches to the nucleus via its proximal end. The molecular machinery connecting the centriole to the nucleus have been termed the head-tail coupling apparatus (HTCA, Gene-Ontology-Term #0120212; (Kierszenbaum et al., 2011).).

Failure to establish or maintain the HTCA leads to the separation of the head and tail (decapitation), and a reduction in fertility. Genetics in humans and mice have identified a number of genes whose loss results in failure at the HTCA including *SUN5, HOOK1, TSGA10, SPATA6, PMFBP1, BRDT, DNAH6, CEP112, PRSS21, OAZ3, CNTROB, IFT88, ODF1, SPATA20, ACTR1,* and *SPATC1L* (reviewed in (Wang et al., 2022) (Graziani et al., 2024). However, the molecular and cellular mechanisms of HTCA assembly, maintenance, and failure in diseased conditions remain largely unexplored.

Much of our limited understanding of the HTCA comes from studies in *Drosophila melanogaster*. Electron microscopy studies have shown that significant rearrangements of the nuclear-centriole attachment occur during spermiogenesis, ultimately resulting in the mature HTCA (Tates, 1971; Tokuyasu, 1975). A number of *Drosophila* proteins, including the SUN domain protein Spag4, the nuclear pore complex, dynein associated proteins, the gravitaxis protein Yuri Gagarin (Yuri), the sytaxin interacting protein Salto, and the centriole protein Pericentrin-like Protein (PLP) localize to the HTCA and disruption of many leads to HTCA failure at some point in spermatid development (Anderson et al., 2009; Augiere et al., 2019; Galletta et al., 2020; Kracklauer et al., 2010; Li et al., 2004; Sitaram et al., 2012; Tates, 1971; Texada et al., 2008). Our lab produced a super-resolution light microscopy map of Spag4 and Yuri rearrangement during HTCA development (Buglak et al., 2024), that discovered an unappreciated role of the atypical centriole, the proximal centriole-like (PCL) structure, in stabilizing the head-tail attachment (Buglak et al., 2024). We showed that the formation of the HTCA occurs in two phases (Figure 1A). The first phase is “Establishment” where the nucleus and the centriole physically come close to one another, which we termed “nuclear search”; the centriole then contacts the nucleus, which we term “attachment”. Both search and attachment require an interplay between microtubules (MTs) emanating from the pericentriolar material at the proximal end of the centriole and the molecular motor dynein localized at the nucleus to ensure the proper orientation of the centriole relative to the nuclear surface (Anderson et al., 2009; Galletta et al., 2020; Li et al., 2004; Sitaram et al., 2012). The second phase of HTCA formation is the “Maintenance” phase, where the centriole “inserts” into the nucleus, then “lateralizes” relative to the long axis of the nucleus (Buglak et al., 2024).

**Figure 1.**
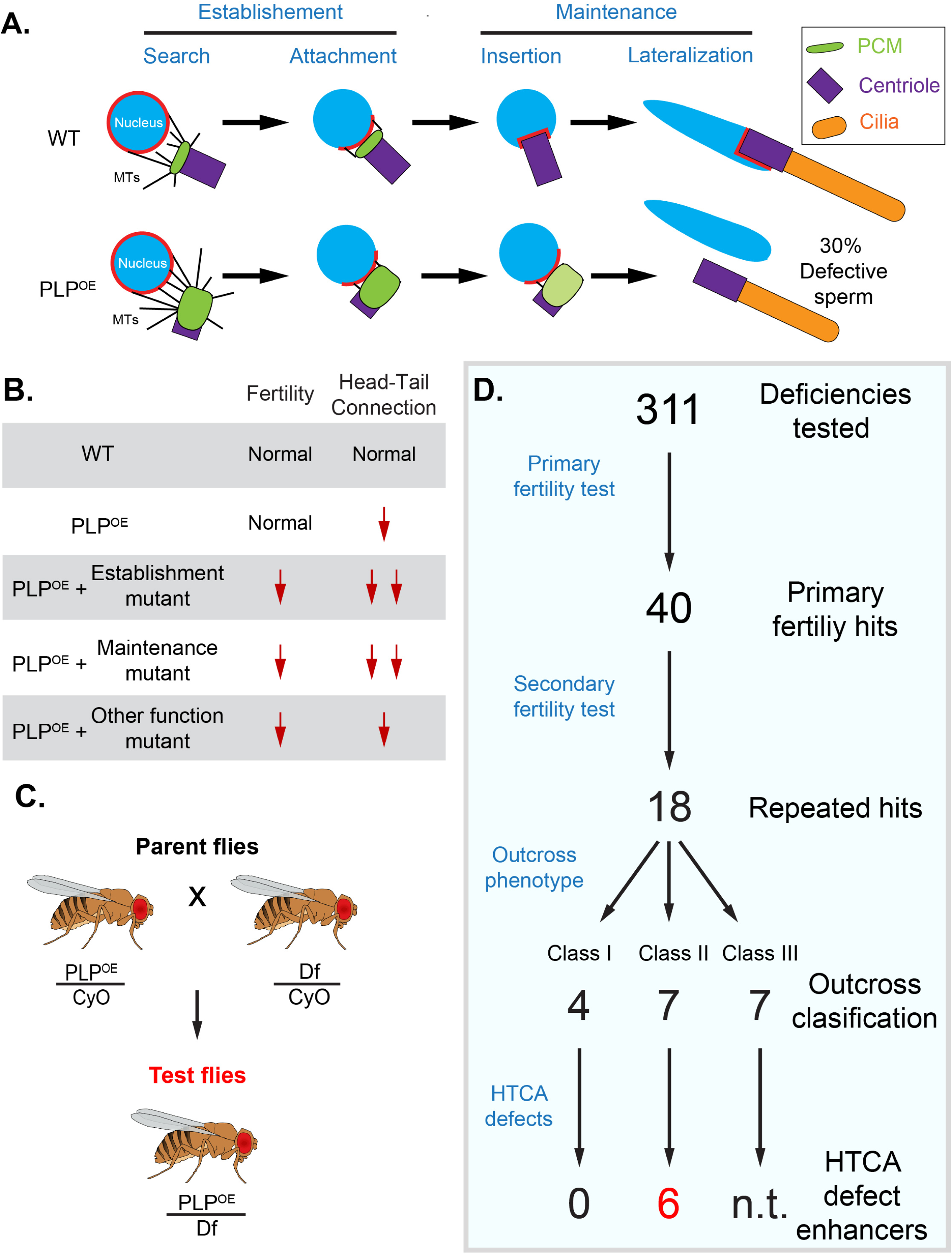
Outline and summary of screen. **A.** Model of the basic stages of HTCA formation during spermiogenesis in wildtype (WT) and PLP^OE^. **B.** Schematic of how the defect in centriole alignment caused by mispositioning of PLP could result in a sensitized background that could be enhanced by heterozygous mutations (+/-) in genes governing other aspects of HTCA formation or spermiogenesis. **C.** Genetic cross used to generate F1 (test) flies. These flies carry ubi-PLP::GFP, which mispositions PLP, and Dfs from the *Drosophila* Df kit. This example shows a Df on the 2^nd^ chromosome. A similar cross was used to obtain F1 flies carrying ubi-PLP::GFP and 3^rd^ chromosome Dfs. **D.** Flow chart of the screen. Experiments are on the left. Results are on the right including the number of Dfs identified at various stages of the screen. The number of F1 males used in each experiment can be found in File S1, Sheet 2.

Spermatogenesis is a linear series of events, which presents challenges to traditional loss of function studies. Classic mutant analysis may only reveal the role of genes in the earliest stages of spermatogenesis masking potential critical roles in spermiogenesis and fertility, the latest stages of spermatogenesis. To overcome this obstacle and identify new genomic regions, and possibly novel genes, required for HTCA functions, we designed and executed a dominant enhancer screen. This type of screen utilizes a sensitized background where a developmental process is already somewhat disrupted, such as loss of a non-essential gene. The hypothesis is that loss of a single copy of another gene (or a set of genes) within the sensitized background will reveal or enhance a measurable phenotype. One advantage of this heterozygous approach is the possibility of revealing phenotypes at later developmental stages that would otherwise be masked by a homozygous mutant phenotype that effected a process at an early developmental stage (reviewed in (St Johnston, 2002)). We chose to pursue this strategy using a previously characterized sensitized background resulting from misexpressing the centrosome protein PLP, which results in centriole alignment defects during HTCA formation and a reduction of ∼30% in functional sperm (Galletta et al., 2020). However, these PLP misexpression flies are fully fertile. Thus, we utilized these flies as a “sensitized” background and combined them with chromosomal deletions from the *Drosophila* deficiency kit. We then screened for a reduction in fertility and an increase in HTCA failure to identify regions of the genome that influence HTCA assembly or maintenance.

## Results and Discussion

Our interest in the sperm head-tail connection motivated our search for regions of the genome that contain new components or effectors of the HTCA. To identify these regions we performed a genome wide deficiency (Df) modifier screen. We reasoned that a semi penetrant HTCA phenotype could be used as a sensitized genetic background to perform this screen. The genetic background we chose was our previously characterized *Drosophila* line that overexpresses the protein PLP (ubi-PLP::GFP), herein referred to as PLP^OE^ (Figure 1A, (Galletta et al., 2014)). This PLP^OE^ background results in significant sperm decapitation and roughly 30% non-functional sperm; however the remaining sperm appear unaffected, the stock is viable, and it does not show a reduction in fertility ((Galletta et al., 2020); Figure S1). We hypothesized that the deletion of a single copy of regions of the genome would enhance the PLP^OE^ phenotype and result in a measurable reduction in fertility (Figure 1B).

To allow for rapid screening of a large swath of the genome, we utilized the *Drosophila* deficiency (Df) kit, which contains a set of overlapping deletions, in total removing almost all coding genes in the fly genome (Cook et al., 2012). We designed a multistage, F1 genetic enhancer screen focusing on the autosomes, chromosomes II and III. To generate ‘Test’ males, PLP^OE^ flies were crossed to flies harboring each of the Dfs on chromosome II and III (Figure 1C, Methods). The F1 males carrying the PLP^OE^ transgene and one copy of the Df were collected and used in testing, beginning with a screen for fertility defects (Figure 1D). Genomic regions identified as hits were subject to secondary screening, including more thorough fertility testing and immunofluorescence microscopy to identify defects in the head-tail connection (Figure 1D).

### Fertility screen and Classifications

The primary screen used male fertility as a readout to identify enhancers. Three F1 test males were individually crossed to two females and the number of progeny was determined (Methods). In total, 311 of the 375 chromosome II and III Dfs (File S1, Sheet 1) were tested for fertility. This primary screen identified 40 Dfs that resulted in a 35% or more reduction in average fertility relative to controls (Figure 1D; Figure 2, colored bars; Methods; File S1, Sheet 2). Given that the initial screen was performed with only 3 F1 males, a secondary screen for fertility was performed using larger numbers of F1 males generated from the 40 primary hits. We found 18 Df lines repeated the results of the primary screen by exhibiting reduced fertility (Figure 1D, Figure 2, cyan bars, File S1, Sheet2).

**Figure 2.**
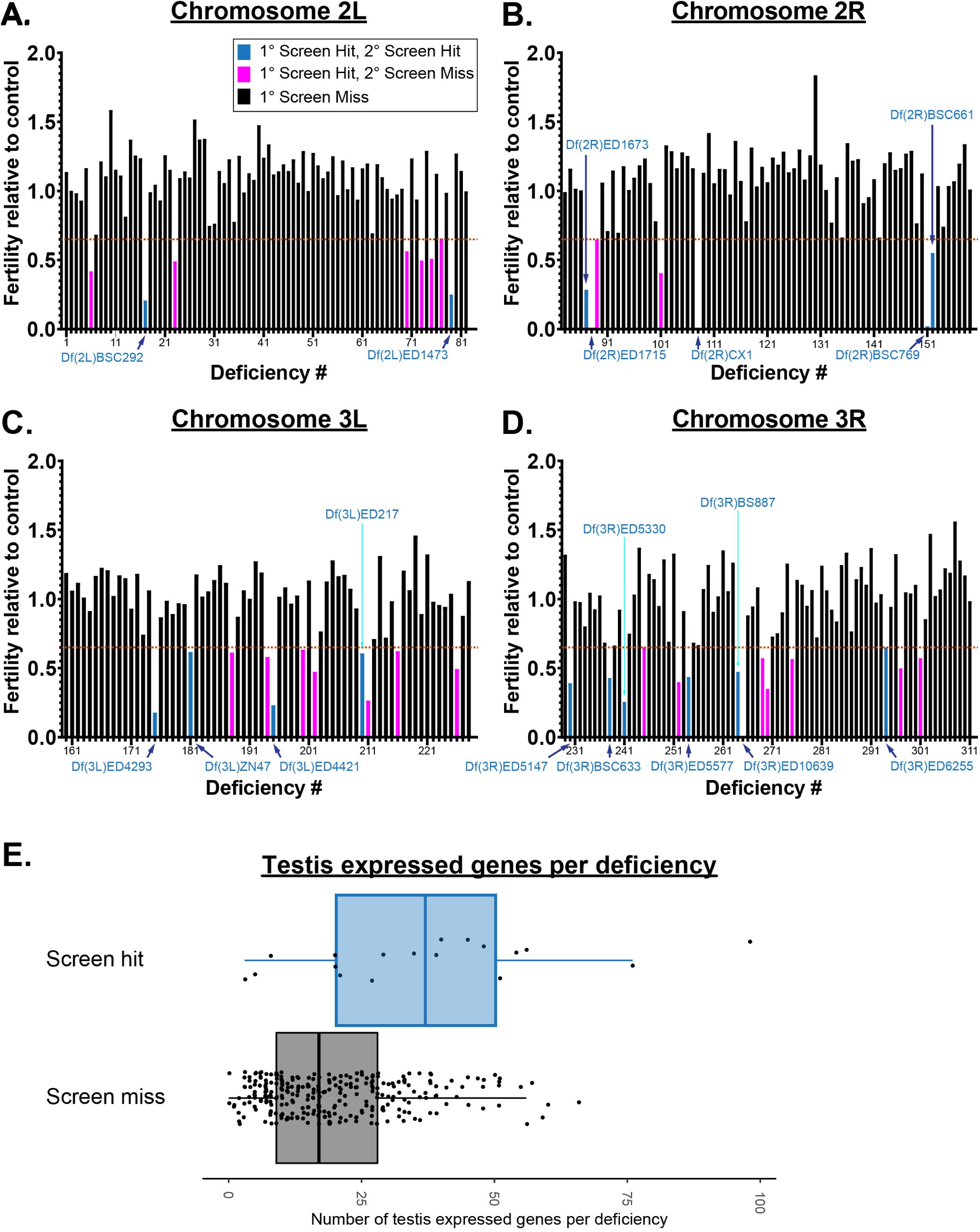
Results of fertility screens of F1 males for chromosome II and III deficiencies. Each bar represents the average fertility, relative to control, determined for F1 flies carrying ubi-PLP::GFP and one of the tested Dfs. The identity of the Df can be determined using the number on the X-axis and the information in File S1, Sheet 1. Dfs are arranged based on the position of the Df along each chromone arm; 2L (**A**), 2R (**B**), 3L (**C**), 3R (**D**). Dashed lines indicate 0.65 fertility relative to control flies. Black bars represent Dfs that did not reduce fertility. Magenta bars represent Df that showed average reduced fertility in the primary screen, but did not show reduced fertility when repeated during the secondary screen. Cyan bars are Df which show reduced fertility in both primary and secondary screens. Df names are indicated in blue text. **E.** The number of testis expressed genes (moderate expression or greater, FPKM ≥ 11) in Dfs that showed a genetic interaction in fertility assays (screen hits) vs those that did not (screen miss). Box represents the interquartile range (IQR) with the line indicating the median. Whiskers extend to the largest or smallest data point or to a distance of 1.5 times the IQR. All data points are shown.

We next determined if the 18 enhancer deficiencies were enriched for genes with testes specific expression. We used FlyAtlas2 and gated our search for genes ‘moderately’ or greater expressed (FPKM >11) in testes. We found that 15 of the 18 Df hits contained a higher number of testis expressed genes than the median found in Dfs that did not genetically interact with PLP^OE^ (Figure 2E). It is important to note that the number of testis expressed genes in a given Df does not predict a genetic interaction as many non-enhancing Dfs in our screen contained a significant number of testes genes (Figure 2E). While it is possible that the 18 Dfs contain genes that have a specific effect on the HTCA, it is equally possible that these Dfs enhanced the ‘reduced fertility’ of PLP^OE^ as a result of loss of several genes with compounding negative effects.

Subsequent analysis of the 18 Dfs that emerged from the secondary screen allowed us to place them into three classifications (Figure 1D, Figure 3). **Class I** included four Dfs that result in reduced fertility only when combined with PLP^OE^ (Figure 3A, B). **Class II** included 7 Dfs that alone have some effect on fertility as heterozygotes, but the combination of the Df and PLP^OE^ result in further reduction in fertility (Figure 3A, C). **Class III** include 7 Dfs that alone have some effect on fertility as heterozygotes, but fertility is similar or improved when combined with PLP^OE^ (Figure 3A, D). Thus, Class III deletions cause a reduction in fertility due to haploinsuficiny in this region with respect to fertility and not due to genetic interaction with PLP^OE^. Interestingly, some of the Dfs in Class III were practically infertile as heterozygotes after outcrossing. This suggests that some genetic feature, like a suppressor mutation or duplication, present in the stocks of these Dfs compensates for the loss of fertility caused by these deletions. Given that we were primarily interested in enhancers of the PLP^OE^ phenotype, we did not consider Class III for further analysis. Thus, the remaining 11 Dfs representing regions of the genome that enhanced the PLP^OE^ fertility defect, each of which have the potential of containing genes that play a role in sperm head-tail connection.

**Figure 3.**
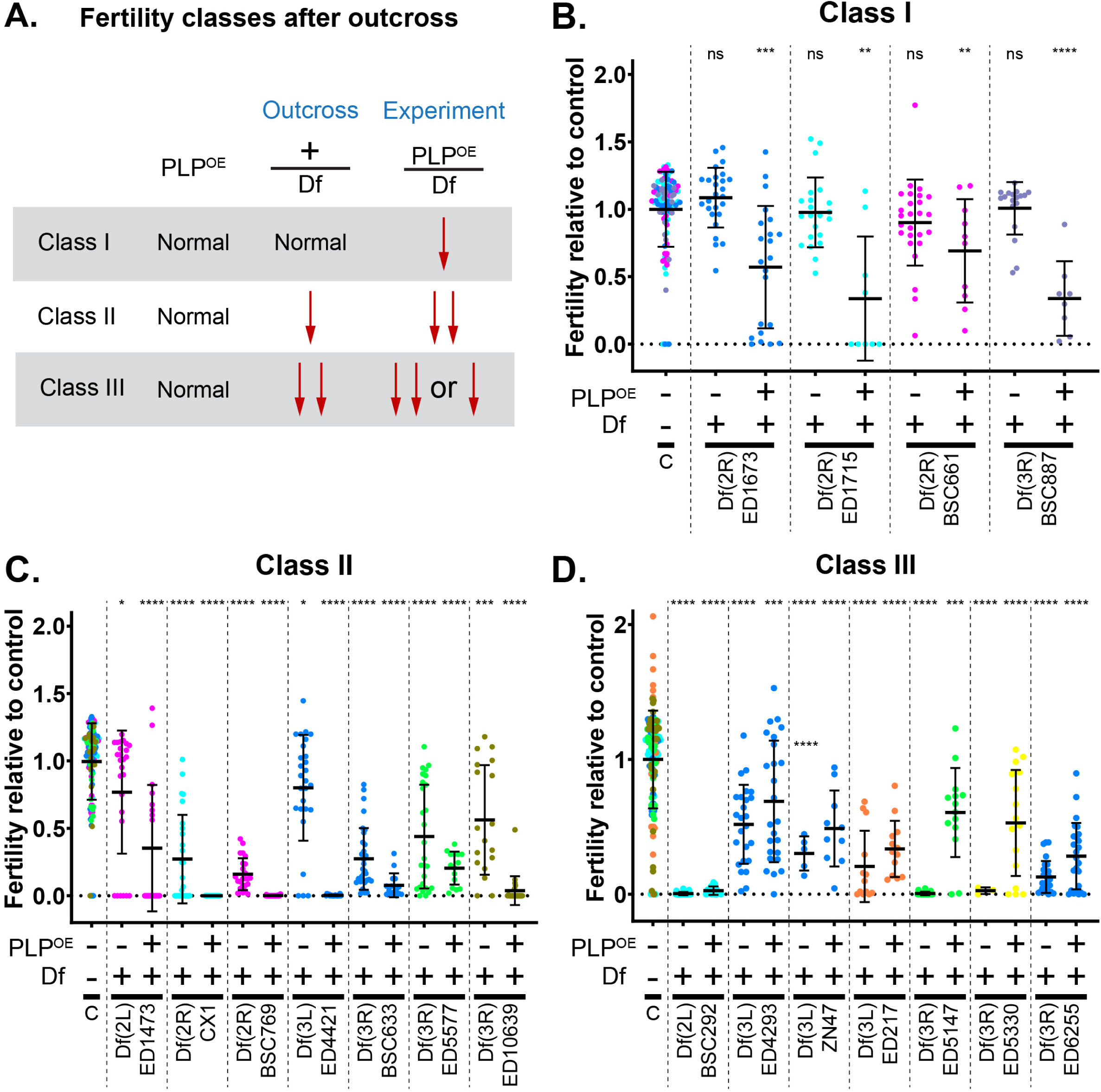
Classification of Fertility Screen Hits. **A.** Schematic diagraming the different classes of Dfs identified in the fertility screen. **B-D.** Fertility relative to control. Data are normalized to the average fertility of control males in a given trial (see methods for details). Each data point is color coded to a given trial (independent experiments). Control data in each panel (B, C and D) include all the trials within a given class. Some trials tested Dfs that were classified into separate classes, which resulted in the use of the corresponding control data in multiple panels. Some of data for these trials has been used in Figures 5. **B.** Class I Dfs: These Df do not affect fertility after outcross, but do affect fertility when combined with ubi-PLP::GFP (PLP misexpression). **C.** Class II Dfs: These Dfs affect fertility after outcross, but further reduce fertility when combined with PLP^OE^. **D.** Class III Dfs: These Dfs have an equal or stronger effect on fertility after outcross than they do when combined with PLP^OE^. Bars are mean ± standard deviation. Statistical comparisons are to control flies and are done via t-test with Welch’s correction when appropriate. Details are in File S1, Sheet 3.* p ≤ 0.05; ** p ≤ 0.01; *** p ≤ 0.001; **** p ≤ 0.0001; ns = not significant

### Dfs affecting sperm head-tail connection

Having identified 11 Dfs in Class I and II, we sought to investigate if any of these genomic regions enhanced the head-tail connection defect seen with PLP^OE^. In our previous studies, we found that many of later stage spermatids (canoe and early needle stage) were defective in their head-tail connection (Galletta et al., 2020). At the time of that study we were unable to precisely quantify head-tail failure due to technical imaging difficulties in wholemount tissues. For our screen, however, we were successful in isolating intact cysts from testes, which allowed for reliable quantification of head tail detachments (Methods). We used anti-Asl antibody to mark the centriole adjunct (Blachon et al., 2008; Klebba et al., 2013)), which allowed us to determine the number of nuclei with attached and unattached centrioles. A centriole adjunct that was directly adjacent to the nucleus was scores as “attached” (Methods, Figure 4A). We found that PLP^OE^ cysts had 21.4 ± 11.2% unattached centrioles, compared to 2.1 ± 4.0 % in controls (Figure 4B). Interestingly, this is slightly lower than the percentage of defective sperm following individualization that we observed by EM in our previous study (Galletta et al., 2020). This could indicate that our current assay of the head-tail connection is conservative and does not detecting all failures, or that additional defects arise during the later states of spermiogenesis.

**Figure 4.**
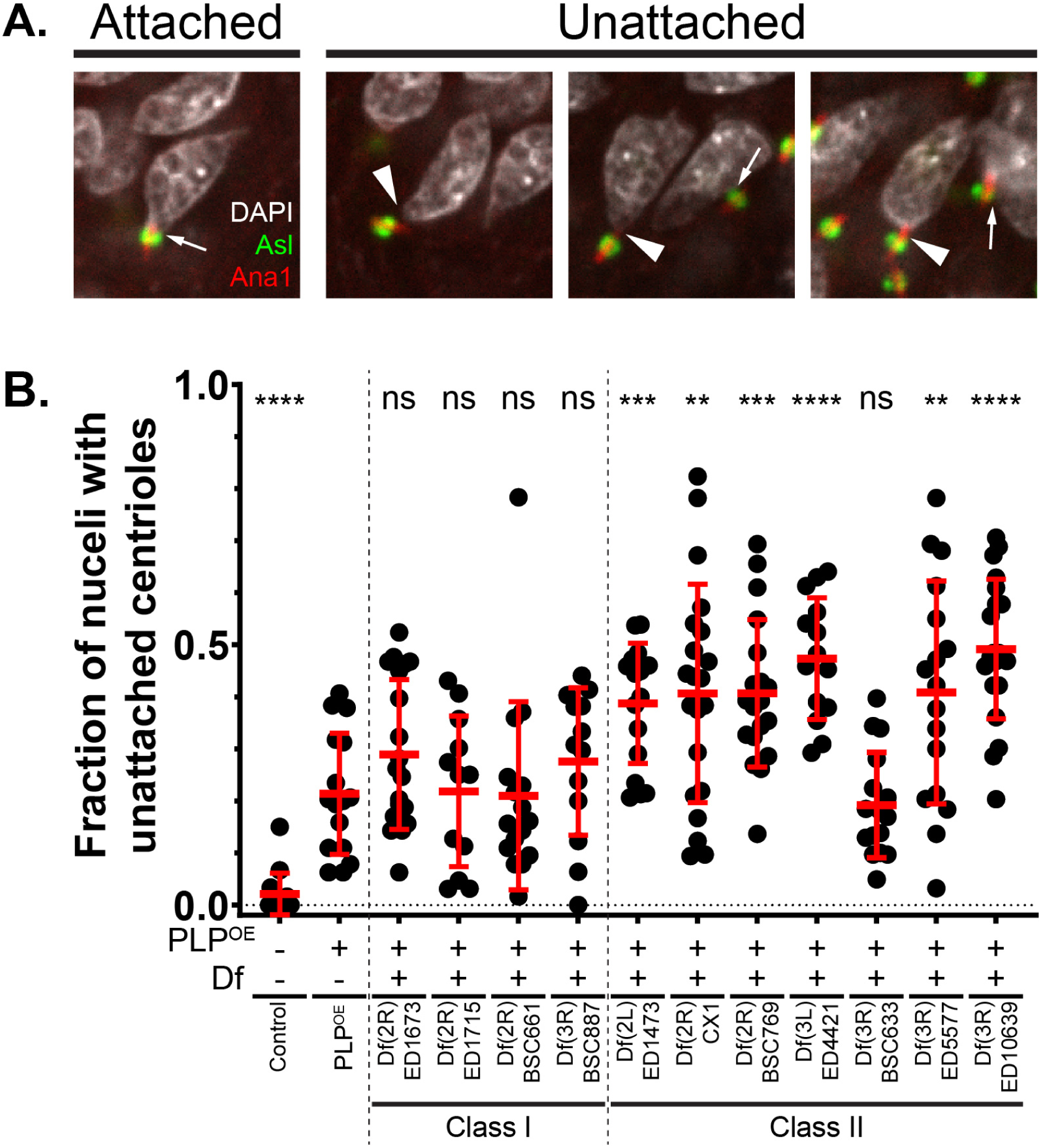
Identifying Dfs that uncover HTCA functions. **A.** Examples of nuclei with attached centrioles (arrows, left) and unattached centrioles (arrowhead, right). DNA (DAPI) is in gray, centriole (Ana1) is in red, centriole adjunct (Asl) is in green. The proximity of the centriole adjunct to the nucleus was used to assess nuclear/centriole attachment. “Attachment” was defined as close apposition of the DAPI signal and the Asl signal. **B.** Fraction of nuclei in cysts with unattached nuclei for flies of the indicated genotypes. Each dot represents a distinct cyst. Bars are mean ± standard deviation. Statistical comparisons are to ubi-PLP::GFP (PLP-misexpression alone) and are done via t-test with Welch’s correction when appropriate. Details are in File S1, Sheet 4. ** p ≤ 0.01; *** p ≤ 0.001; **** p ≤ 0.0001; ns = not significant

By examining F1 males from the 11 Class I and Class II Dfs, we found that none of the class I males enhanced the head-tail connection of PLP^OE^ (Figure 4B, columns 3-6). It is likely that these Dfs cause infertility in combination with PLP^OE^ by affecting a process other than HTCA formation. In contrast, 6 of the 7 class-II Dfs resulted in an increased frequency of HTCA defects compared to PLP^OE^ alone (Figure 4B, columns 7-13), suggesting that these Dfs contain genes that influence HTCA formation/maintenance. Taken together, this screen has identified 11 regions of the genome that contain gene(s) that influence some aspect of male fertility, 6 of which affect HTCA formation.

### Characterization of HTCA screen hits

To begin to understand why these enhancer Dfs genetically interact with PLP^OE^, we aimed to narrow the genomic regions containing the responsible genetic elements. To accomplish this, we obtained smaller Dfs within each region from the Bloomington *Drosophila* Stock Center and tested them for genetic interaction with PLP^OE^ by fertility assay (Figure 5). The simple assumption was that a smaller Df that resulted in the same degree of enhancement is likely to contain a genetic element with a sufficient enough interaction to have been the causative element in the initial screen. In contrast, smaller Dfs resulting in no enhancement would not contain a gene of interest. In a cases where none of the smaller Dfs result in as strong of an enhanced phenotype, we hypothesized that loss of two or more genes within the larger Df are responsible for enhancing PLP^OE^ phenotypes. These assumptions will help guide future work aimed at understanding the specific genes and mechanisms that contribute to these genetic interactions.

**Figure 5.**
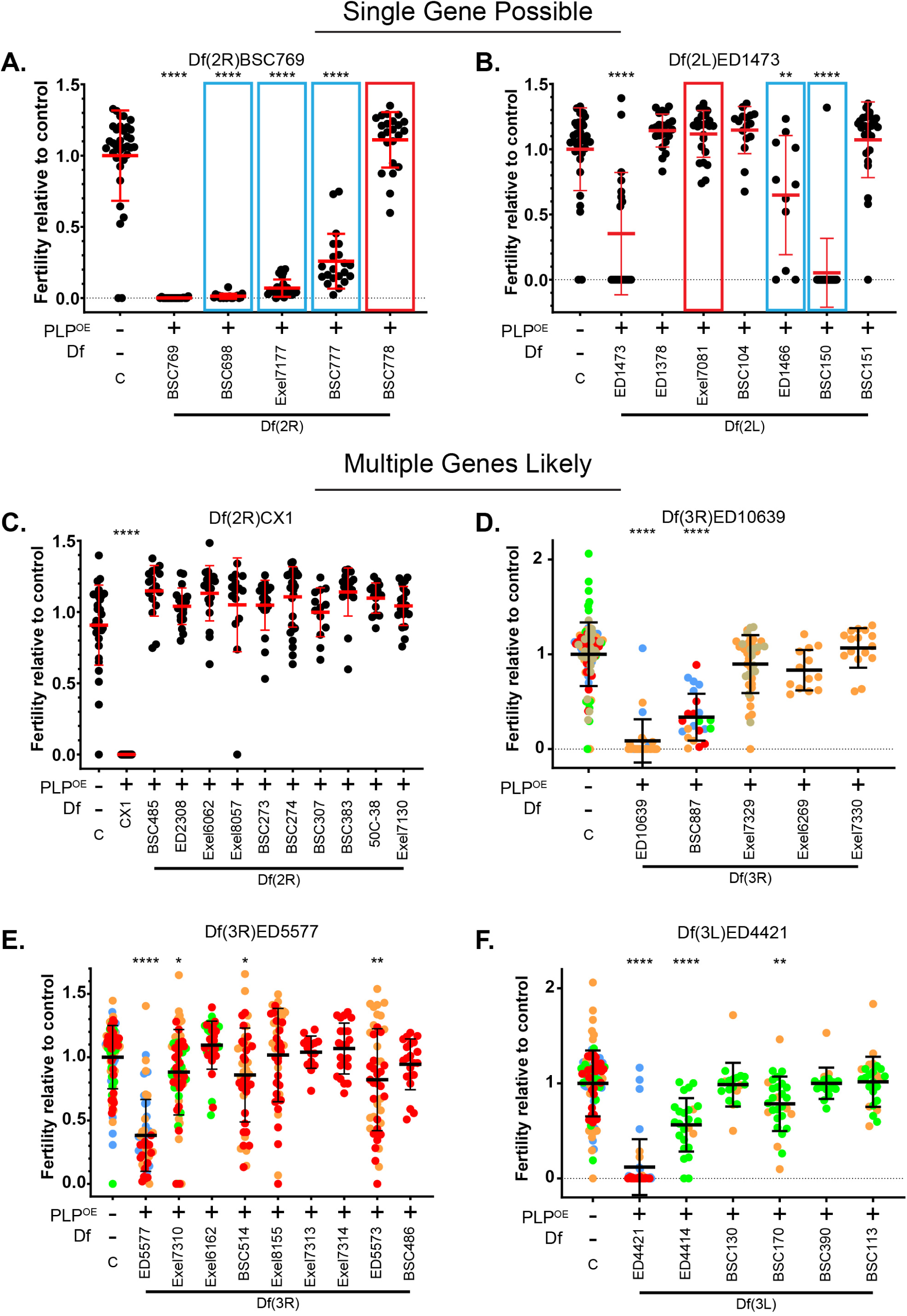
Using smaller Dfs to refine the region of interest. Fertility tests of smaller Dfs, relative to control, that subdivide the region of the genome uncovered by fertility screen hits that showed an enhancement of the HTCA defect of PLP^OE^. **A-B.** “Single Gene Possible” Category. Smaller deletions within these regions showed comparable phenotypes to the original screen hit Df (blue boxes). Deletions that did not show an genetic interaction, but were used to narrow the likely region of interested are boxed in red. **C-F.** “Multiple Gene Likely” category. No smaller Df in this region was sufficient to reproduce the degree of enhancement seen with the original screen hit, suggesting multiple loci within the original Df, all of which are not disrupted in any smaller Df, need to be disrupted to show the original degree of enhancement. Statistical comparisons are to control flies and are done via t-test with Welch’s correction when appropriate. Details are in File S1, sheet 5. *≤ 0.05 p ** p ≤ 0.01; *** p ≤ 0.001; **** p ≤ 0.0001. In cases where multiple fertility tests were performed independently each data point is color coded to indicate various trials. Some of the data in this figure is reproduced from Figure 3 and some experiments performed simultaneously share control data.

We performed fertility assays on the smaller Dfs and sorted them into two categories. The first category is “Single Gene Possible” where our analysis of the genetic interaction with smaller Df(s) enhanced the PLP^OE^ fertility defect at a similar level as the larger Df from the original screen. In these cases we hypothesize that a single gene within each smaller Df is sufficient to cause the enhancement. Within the 6 original Dfs, we found two “Single Gene Possible” Dfs – Df(2R)BSC769 and Df(3R)ED1473 (Figure 5A, B). The second category is “Multiple Genes Likely” where no smaller Df caused a genetic interaction equal to the larger Df from the original screen. In these cases we hypothesized that some combination of multiple genes/regions are responsible for the level of enhancement measured in the original screen. Within the 6 original Dfs, we found four “Multiple Genes Likely” Dfs – Df(2R)CX1, Df(3R)ED10639, Df3R)ED5577, and Df(3L)ED4421 (Figure 5C-F).

### Using gene expression data to investigate possible functions and prioritize screen hits

To begin to explore the genetic landscape within the original 6 screen Dfs, we used the results of fertility tests of smaller Dfs (Figure 5) to define the “most likely” regions. We then used FlyAtlas 2 to map the position of all genes with “moderately expressed” testes transcript levels (FPKM ≥ 11) or higher (Gillen, A. 2023.11.14, FlyAtlas2 2023 Data, FBrf0258027; (Krause et al., 2022; Leader et al., 2018) in these regions. For each of the “most likely” regions identified (gray shaded regions, Figure 6A, 7A, Figure S2A, S3A, S4A, S5A), we examined: 1) The level of expression of RNAs in testes and other tissues from male flies using data from FlyAtlas 2 (Gillen, A. 2023.11.14, FlyAtlas2 2023 Data, FBrf0258027; (Krause et al., 2022; Leader et al., 2018). 2) The presence of the protein product in the most recent *Drosophila* sperm proteome (Garlovsky et al., 2022). 3) The relative expression of these genes in the developing germline and somatic cysts using single nucleus RNAseq (snRNAseq) data from Fly Cell Atlas (Li et al., 2022). Below we describe some of the features of the “most likely” regions within both the “Single Gene Possible” and “Multiple Genes Likely” categories.

**Figure 6.**
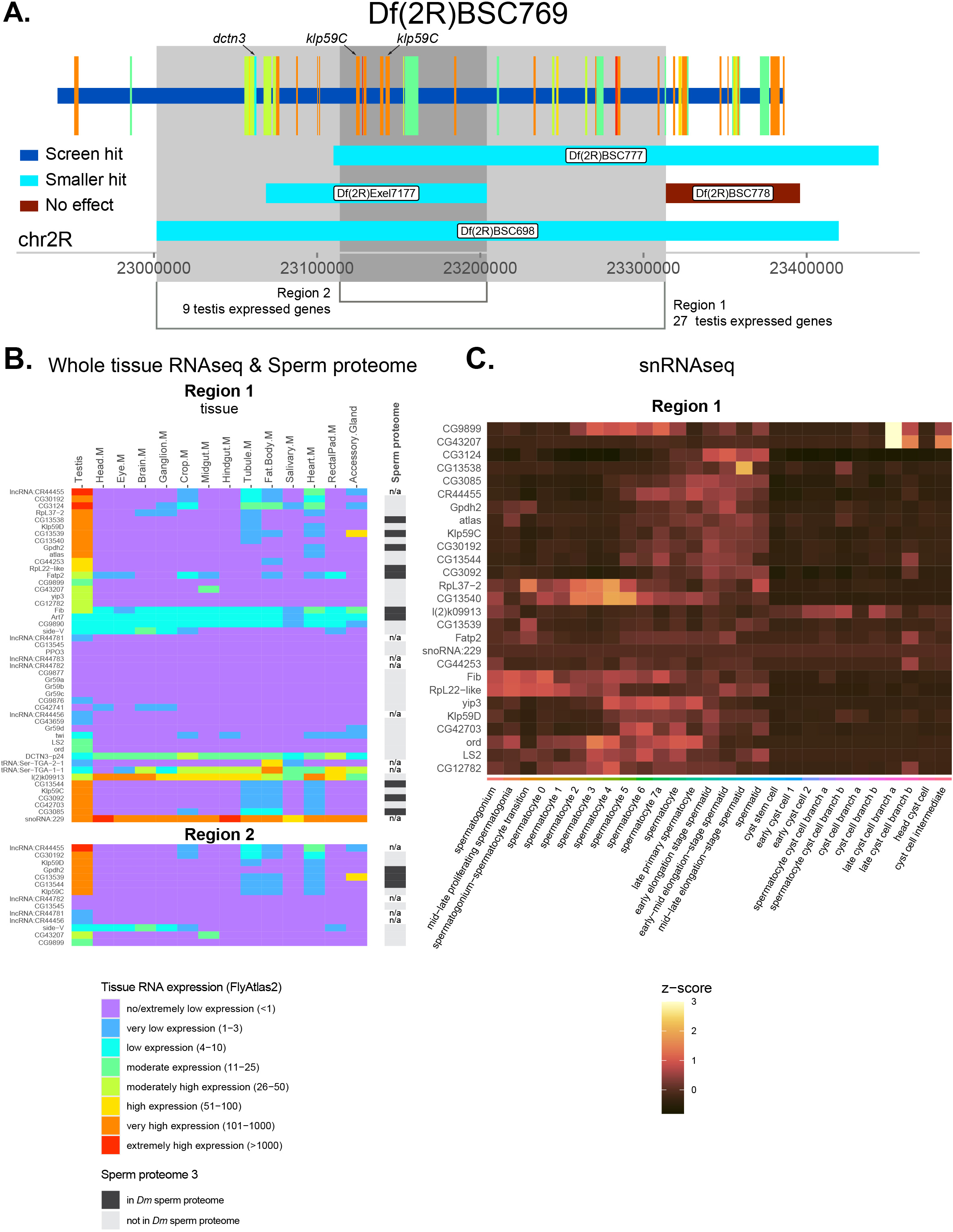
Analysis of screen hit Df(2R)BSC769. **A.** Genomic region of DF(2R)BSC769. Horizontal bars represent regions of the genome removed by Df screen hits and each smaller Df tested. Interaction status color code in inset. Genes removed by the Df from the screen are represented by vertical bars with their width indicating the region of the genome they occupy. Gene color indicates expression level (FPKM) in testis from FlyAtlas 2 (legend in B). Except for *dctn3* only genes with moderate or higher expression are displayed. Axis is FlyBase Gene Coordinates. Grey regions represent the regions of the genome discussed in the Results/Discussion and the specific genes discussed are indicated. **B.** Heatmap of the FPKM from RNAseq (FlyAtlas 2) of genes in grey region in A. for various tissues. Rightmost column indicates the presence or absence of proteins encoded by these genes in the sperm proteome 3. **C.** Heatmap of snRNAseq data from FlyCellAtlas of gens in C. with moderate or higher expression in the testis. **A.** Genomic region of DF(2R)CX1. Horizontal bars represent regions of the genome removed by Df screen hit and each smaller Df tested. Interaction status color code in inset. Genes removed by the Df from the screen are represented by vertical bars with their width indicating the region of the genome they occupy. Gene color indicates expression level (FPKM) in testis from FlyAtlas 2 (legend in D). Axis is FlyBase Gene Coordinates. Grey region represents the region of the genome discussed in the Results/Discussion where no smaller deletion was available and the specific genes discussed are indicated. **B.** Heatmap of the FPKM from RNAseq (FlyAtlas 2) of genes in grey region in A. for various tissues. Rightmost column indicates presence or absence of proteins encoded by these genes in the sperm proteome 3. **C.** Heatmap of snRNAseq data from FlyCellAtlas of gens in C. with moderate or higher expression in the testis.

**Figure 7.**
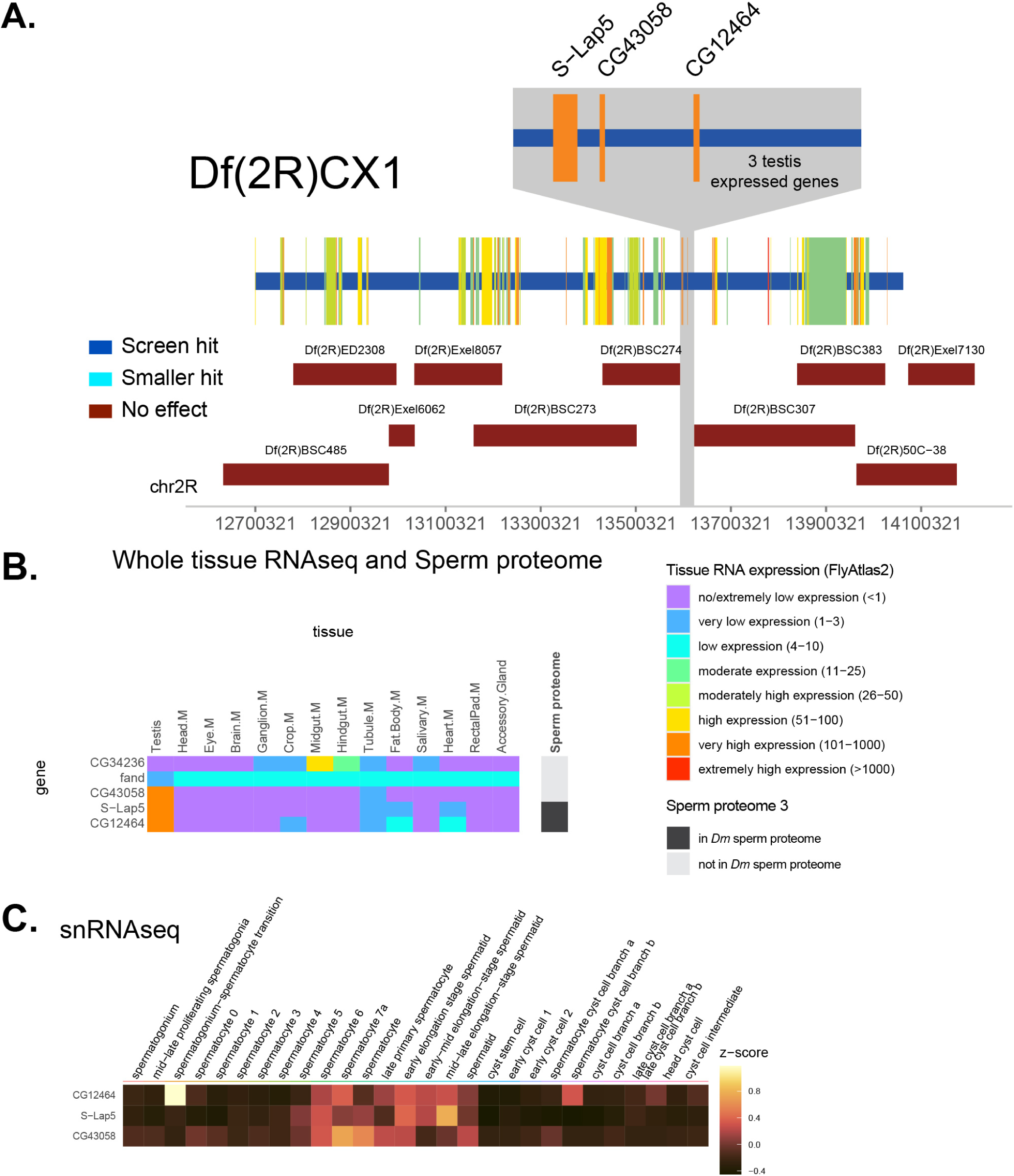
Analysis of screen hit Df(2R)CX1. **A.** Genomic region of DF(2R)CX1. Horizontal bars represent regions of the genome removed by Df screen hit and each smaller Df tested. Interaction status color code in inset. Genes removed by the Df from the screen are represented by vertical bars with their width indicating the region of the genome they occupy. Gene color indicates expression level (FPKM) in testis from FlyAtlas 2 (legend in D). Axis is FlyBase Gene Coordinates. Grey region represents the region of the genome discussed in the Results/Discussion where no smaller deletion was available and the specific genes discussed are indicated. **B.** Heatmap of the FPKM from RNAseq (FlyAtlas 2) of genes in grey region in A. for various tissues. Rightmost column indicates presence or absence of proteins encoded by these genes in the sperm proteome 3. C. Heatmap of snRNAseq data from FlyCellAtlas of gens in C. with moderate or higher expression in the testis.

### “Single Gene Possible” Deficiencies

#### Df(2R)BSC769

The region of the genome removed by Df(2R)BSC769 contains 85 genes, 45 of which are expressed at moderate or higher levels in the testis (Figure 6A). Three of the tested smaller Dfs in this region genetically interacted with PLP^OE^ in fertility assays and all to a similar degree as the Df from the screen (Figure 5A). Based on these results we hypothesize that each of Df(2R)BSC698, Df(2R)Exel7177, and Df(2R)BSC777 could contain a single gene responsible for the phenotype observed from the larger original screen Df (Df(2R)BSC769). Therefore, we examined genes within Df(2R)BSC769 (Figure 6A, Region 1) that overlapped with the smaller, positive-hit Dfs (Figure 5A blue box; Figure 6A, blue bars), minus the region Df(2R)BSC778 that did not enhance PLP^OE^ (Figure 5A red box; Figure 6A, red bars). We also gave added focus to a narrower region where all three of the smaller, positive-hit Dfs overlapped (Figure 6A, Region 2). Both region 1 (48 genes) and 2 (14 genes) stand out for the significant number of genes that have “very-high” or “extremely-high” expression in the testes (Figure 6B), while also showing nearly testis specific expression. Furthermore, analysis of snRNAseq data revealed that many of the testis expressed genes are enriched in the germline cells of the testis (Figure 6C), consistent with possible direct roles in spermatogenesis. Interestingly, Region 1 includes 10 genes encoding proteins found in the sperm proteome, 3 of which are also in Region 2 (Gpdh2, CG13539, CG13544). The presence of genes that encode proteins found in the mature sperm could be indicative of roles in the mature sperm that might manifest later in spermiogenesis or in the final product.

While there are many candidate genes in this region that might contribute to interaction with PLP^OE^, we highlight three genes which directly relate to microtubule dynamics and microtubule based movement, processes critical for HTCA formation and highly related to the phenotype caused by PLP^OE^ (Anderson et al., 2009; Galletta et al., 2020; Li et al., 2004; Sitaram et al., 2012). Two of these genes are encode microtubule depolymerizing kinesin-13 orthologs, Klp59C and Klp59D (Figure 6A,B,C). These genes are of great interest because microtubules are present around the site of HTCA formation throughout spermiogenesis and their organization changes throughout ((Anderson, 1967; Riparbelli et al., 2020).

These changes would require that the microtubule cytoskeleton remodel, a process known to involve depolymerization of microtubule polymers, possibly via the Klp59C and Klp59D. The third gene encodes a component of the Dynein/Dynactin complex, DCTN3/p24. While this is not expressed highly in the testis, it is a gene is of particular interest given the known role of dynein and dynactin HTCA formation (Anderson et al., 2009; Li et al., 2004; Sitaram et al., 2012). We hypothesize that DCTN3/p24 serves as a critical regulator of Dynein/Dynactin to ensure proper spatial and temporal function of this ubiquitous motor. Taken together, a reasonable hypothesis for future testing is that the original Df(2R)BSC769 may have interacted because it removes microtubule motors that could be regulating the pull of the centriole towards the nucleus and / or the dynamics and remodeling of the microtubule array during spermiogenesis.

#### Df(2L)ED1473

This deletion removes 178 genes, 54 of which are expressed at moderate or higher levels in the testes (Figure S2A). Two Dfs, Df(2L)ED1466 and Df(2L)BSC150, of the six smaller Dfs tested in this region showed an interaction with PLP^OE^ in fertility assays similar to or stronger than Df(2L)ED1473 (Figure 5B, blue boxes), excluding the region removed by Df(2L)Exel7081, which did not interact (Figure 5B, red boxes), and the region beyond the end of Df(2L)ED1473. We therefore examined all of the genes present in the region covered by both these Dfs, and excluded genes (Figure 5B, S2A). This region contains a total of 10 genes, 6 (5 protein coding genes) of which are expressed in the testis at moderate or high levels, and 4 are present in the sperm proteome (Figure S2B). All of these genes have some enrichment in the germline (Figure S2C). Of the coding genes, two encode known proteins, Steppke (Step) an Arf-guanine-nucleotide exchange factor (Fuss et al., 2006), that has been shown to have a role in germ cell segregation in the early embryo (Lee et al., 2015), and Eukaryotic translation elongation factor 2 (eEF2; (Grinblat et al., 1989). The remaining genes, *CG1416*, *CG2225* and *CG11630* are uncharacterized genes with little to no insight into their functions (Flybase).

### “Multiple Genes Likely”

#### Df(2R)CX1

The region of the genome removed by this deletion contained 214 genes, 98 of which were expressed at moderate or higher levels in the testis, more than any other Df tested in our screen. Interestingly, none of the 10 smaller Dfs resulted in a genetic interaction with PLP^OE^ (Figure 5C, 7A), thus falling in the “multiple genes likely” category. Our primary hypothesis is that more than one gene within Df(2R)CX1must be removed to lower fecundity in the PLP^OE^ background. However, there is a small region that was not uncovered by any of the tested Dfs (Figure 7A). This region contains only 5 genes, 3 of which (*CG12464*, *CG43058*, *s-lap5*) have “very high expression” in testes (Figure 7B), and snRNAseq data indicates that expression of these 3 genes is enriched in the germline (Figure 7C). Furthermore, these 3 genes have almost no expression in other tissues and the protein products of 2 of these 3 are found in the sperm proteome (Figure 7B). CG12464 is predicted to be located in mitochondria as part of respiratory chain complex I (Flybase), and thus unlikely to be involved in enhancing the HTCA defect of PLP^OE^. *CG43058* encodes a RING-type Zinc Finger and may encode an E3 ubuiquitn-protein ligase (Flybase). One hypothesis is that CG43058 directly or directly influences PLP positioning on the centriole and thus further enhances the PLP^OE^, which would be similar to our previous work showing that regulation of PLP protein levels by protein degradation is critical for its proper function in head-tail connection (Galletta et al., 2023) . CG43058 was not one of the proteins tested in our previous study, and therefore remains a candidate PLP regulator. Finally, the third gene with high testes expression in this small region is *sperm-leucylaminopeptidase* 5 (*s-lap5*), which is predicted to function in mitochondria, and play a role in the migration of investment cones during sperm individualization (Laurinyecz et al., 2019). Interestingly, failure of sperm individualization is associated with defects in the head-tail connection of the sperm (Fabian and Brill, 2012; Galletta et al., 2020; Kracklauer et al., 2010; Texada et al., 2008), raising the possibility of a function of S-Lap5 in the HTCA.

#### Df(3R)ED10639

This deletion removed 48 genes, 20 that are expressed at “moderate” or higher levels in the testes (Figure S3A). Of the 4 tested smaller Dfs, only Df(3R)BSC887 showed an interaction (Figure 5D, S3A). The region of overlap between Df(3R)ED10639 and Df(3R)BSC887 that also does not overlapping with the other smaller Dfs (Figure S3A, Region 1), does not contain any protein coding genes that are not also removed by the smaller Dfs. (Figure S3A). It is thus more likely that the lack of a phenotype with the smaller Dfs indicates that the simultaneous loss of multiple loci within Df(3R)ED10639 or Df(3R)BSC887 (Region 2) is requited to genetically enhance PLP^OE^. We therefore placed Df(3R)ED10639 in the “multiple genes likely” category. Region 2 contains several genes that are strongly expressed in the testis (some almost exclusively), and many genes found in the sperm proteome (Figure S3B). The expression of many of these genes is enriched in the germline (Figure S3C). In addition, Region 2 contains the gene *asunder (asun)*, which does not appear to be expressed strongly in the testis RNAseq, but is important for regulating dynactin during HTCA formation (Anderson et al., 2009). Thus, this region contains many genes that could influence HTCA formation.

#### Df(3R)ED5577

The deficiency removes 133 genes, 51 expressed at “moderate” levels or higher in the testis (Figure S4A). Three of the eight smaller deletions in this region interacted with PLP^OE^ in fertility assays (Figure 5E), but none to the same degree as Df(3R)ED5577. Thus this falls in the “multiple gene likely” category. Based on the arrangement of the Dfs, the simplest model points to at least two distinct regions that contribute to fertility (Figure S4A). Region 1 only contains 4 genes, all of which have low or very low expression in the testis in the FlyAtlas2 data and their gene products are not found in the sperm proteome (Figure S4C, Region 1). So we have little insight into how this region contributes to the fertility interactions seen with Df(3R)Exel7310 and Df(3R)BSC514. Region 2 contains 8 genes expressed at moderate or higher levels in the testis (Figure S4C, Region 2), all of which have some enrichment in the germline in the snRNAseq dataset (Figure S4D), but none suggest any obvious hypotheses for function.

#### Df(3L)ED4421

This deficiency removes 89 genes, 39 expressed at moderate or higher levels in the testis (FIgureS5A). Two of the four smaller Dfs in this region interacted with PLP^OE^, although neither showed as severe a phenotype as Df(3L)ED4421 (Figure 5F), making this a “multiple gene likely” category Df. This suggests multiple loci in this region contribute to the phenotype, likely located in two regions. Region 1 contains 8 genes expressed at moderate or higher levels in the testis (Figure S5A,B) and 7 of these appear to be enriched in the germline (Figure S5C). The second region is defined by Df(3L)BSC170. Interestingly, while Df(3L)BSC170 did interact with PLP^OE^, two smaller deletions which together completely overlap with Df(3L)BSC170 did not (Figure 5F, S5A). A simple model is that Df(3L)BSC170 removes 2 (or more) loci that in combination cause an interaction, but neither of the smaller deletions removes both. If this is the case, there are only 3 testis expressed genes that are removed by Df(3L)BSC170 and are also removed by either Df(3L)BSC130 or Df(3L)BSC390, but not both, *arginine kinase 1* (*argk1*), *CG4911* and *unconventional SNARE in the ER 1* (*use1*) (Figure S5A). Only 2 of these, CG4911 and use1 are enriched in the germline relative to the somatic cyst cells (Figure S5C). Interestingly CG4911 is an F-box protein predicted to be involved in protein ubiquitination (Flybase). As noted above, regulation of PLP levels is critical for HTCA formation. CG4911 was not tested in our screen of regulators of PLP degradation (Galletta et al., 2023) and therefore is an interesting candidate for future examination.

## Summary

Genetic enhancer screens are a powerful tool to help identify unknown players in a given process. For most processes, one copy of a gene is sufficient to perform its role. However, in a sensitized background this may not always be the case. This allows one to screen for the effects of loss of a single copy of a gene, even if that gene is essential for development (St Johnston, 2002). Furthermore, such a screen can identify genetic interactions that require the loss of several genes in combination. This appears to be the case for several of the Dfs identified in this screen. Even when our results suggests that a narrow region of the genome is most likely responsible for the interaction, our analysis of RNAseq, snRNAseq and proteomic datasets reveals several likely candidates. Long-term, directed studies will be needed to better understand how these regions of the genome influence HTCA formation and function.

## Data Availability Statement

All relevant data can be found within the article and its supplementary information. The DOI for all raw data is **XXXXXXX (will be provided if and when accepted)**

## Acknowledgments

We thank the Bloomington *Drosophila* stock center for flies; the Xu lab with help with testes dissections members of the Rusan lab for helpful discussion.

## Supplemental Material

### Supplemental Figures

**Figure S1.**
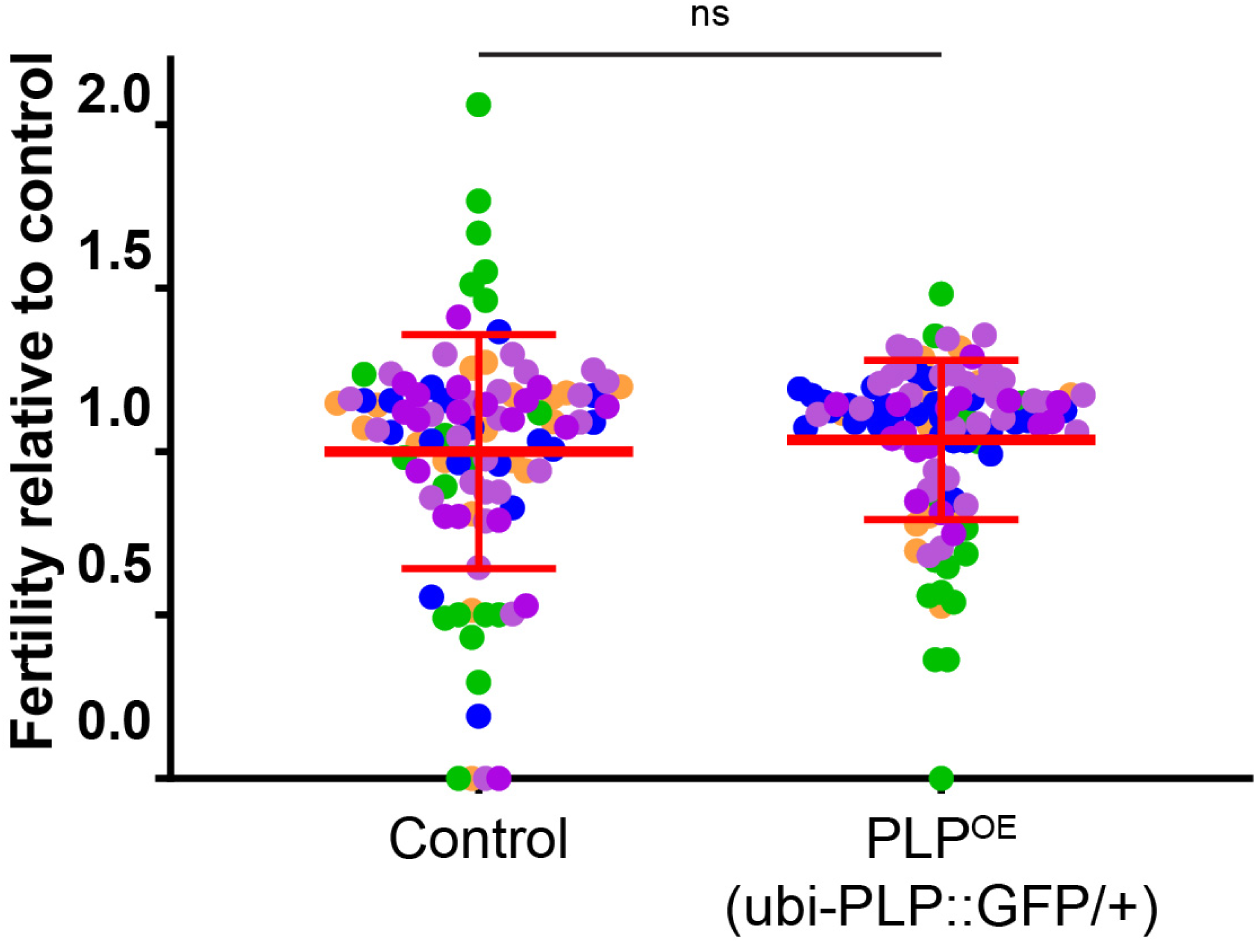
PLP overexpression does not affect male fertility. Male fertility is normal in flies overexpressing PLP::GFP using the ubi promoter from a single transgene (n=98) compared to control (n=102). This is the “sensitized” strain background used in the Df kit screen. Some of the control data has also been used in other figures. Each data point is color coded to a given trial. Statistical comparisons done via t-test with Welch’s correction. ns = not significant

**Figure S2.**
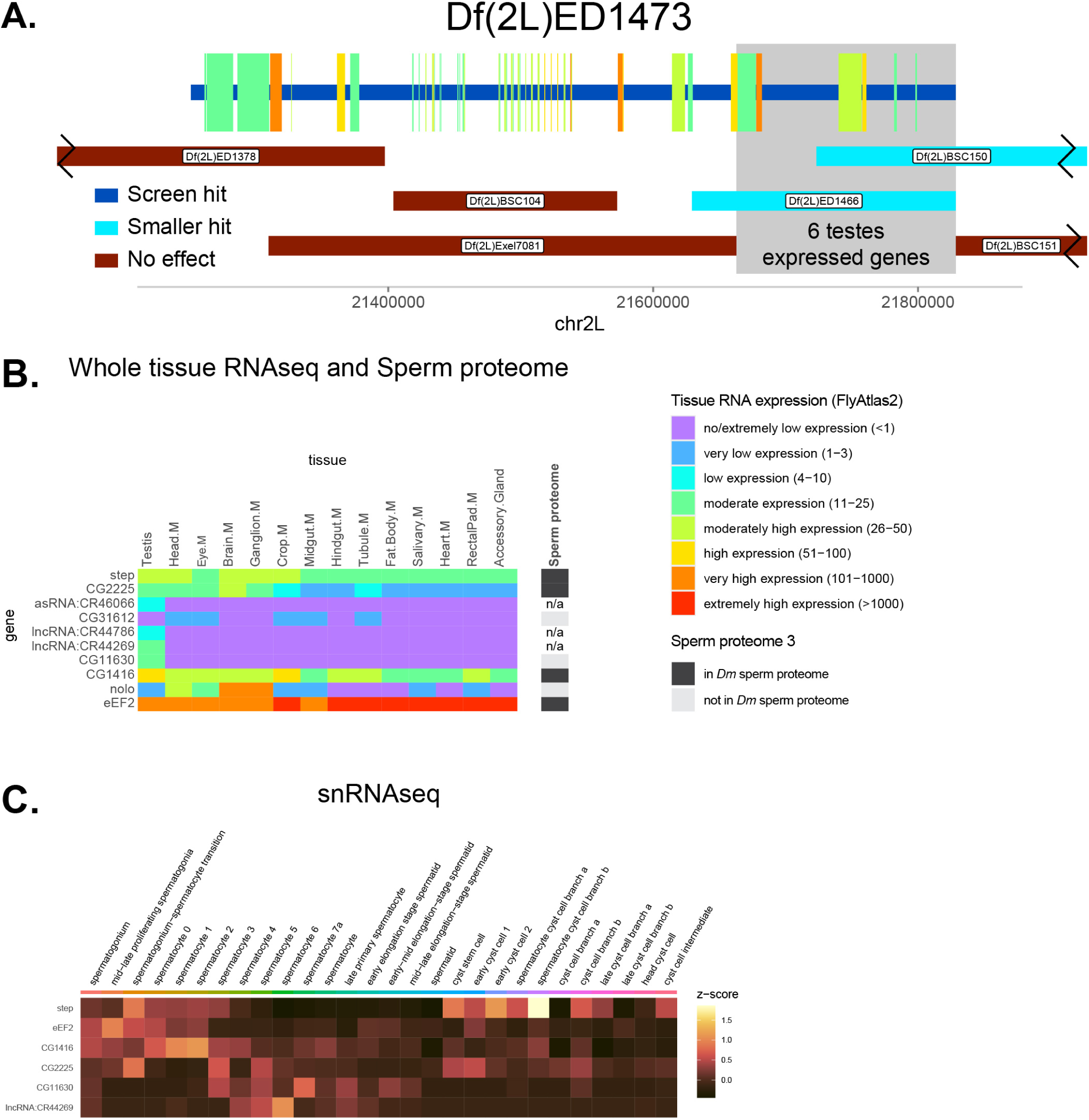
Analysis of screen hit Df(2L)ED1473. **A.** Genomic region of DF(3R)ED1473. Horizontal bars represent regions of the genome removed by Df screen hits and each smaller Df tested. Interaction status color code in inset. Genes removed by the Df from the screen are represented by vertical bars with their width indicating the region of the genome they occupy. Gene color indicates expression level (FPKM) in testis from FlyAtlas 2 (legend in B). Axis is FlyBase Gene Coordinates. Grey region represents the region of the genome discussed in the Results/Discussion. **B.** Heatmap of the FPKM from RNAseq (FlyAtlas 2) of genes in grey region in A. for various tissues. Rightmost column indicates presence or absence of proteins encoded by these genes in the sperm proteome 3. **C.** Heatmap of snRNAseq data from FlyCellAtlas of gens in C. with moderate or higher expression in the testis.

**Figure S3.**
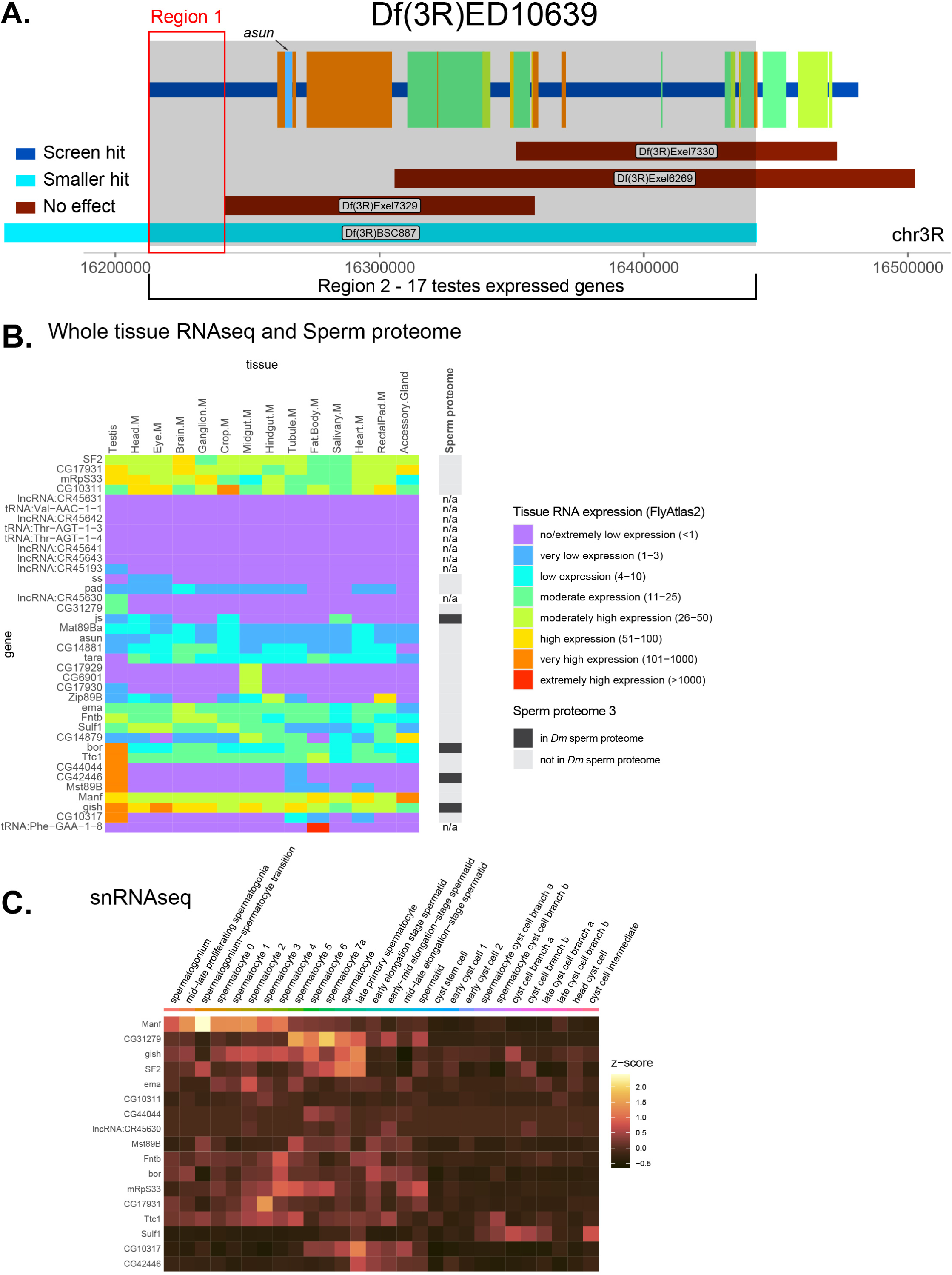
Analysis of screen hit Df(3R)ED10639. **A.** genomic region of DF(3R)ED10639. Horizontal bars represent regions of the genome removed by Df screen hits and each smaller Df tested. Interaction status color code in insets. Genes removed by the Df from the screen are represented by vertical bars with their width indicating the region of the genome they occupy. Gene color indicates expression level (FPKM) in testis from FlyAtlas 2 (legend in B). Except for *asun* only genes with moderate or higher expression are displayed. Axis is FlyBase Gene Coordinates. Regions of the genome discussed in the Results/Discussion are indicated. **C.** Heatmap of the FPKM from RNAseq (FlyAtlas 2) of genes in grey region in A. for various tissues. Rightmost column contains indicates presence or absence of proteins encoded by these genes in the sperm proteome 3. **C.** Heatmap of snRNAseq data from FlyCellAtlas of gens in C. with moderate or higher expression in the testis.

**Figure S4.**
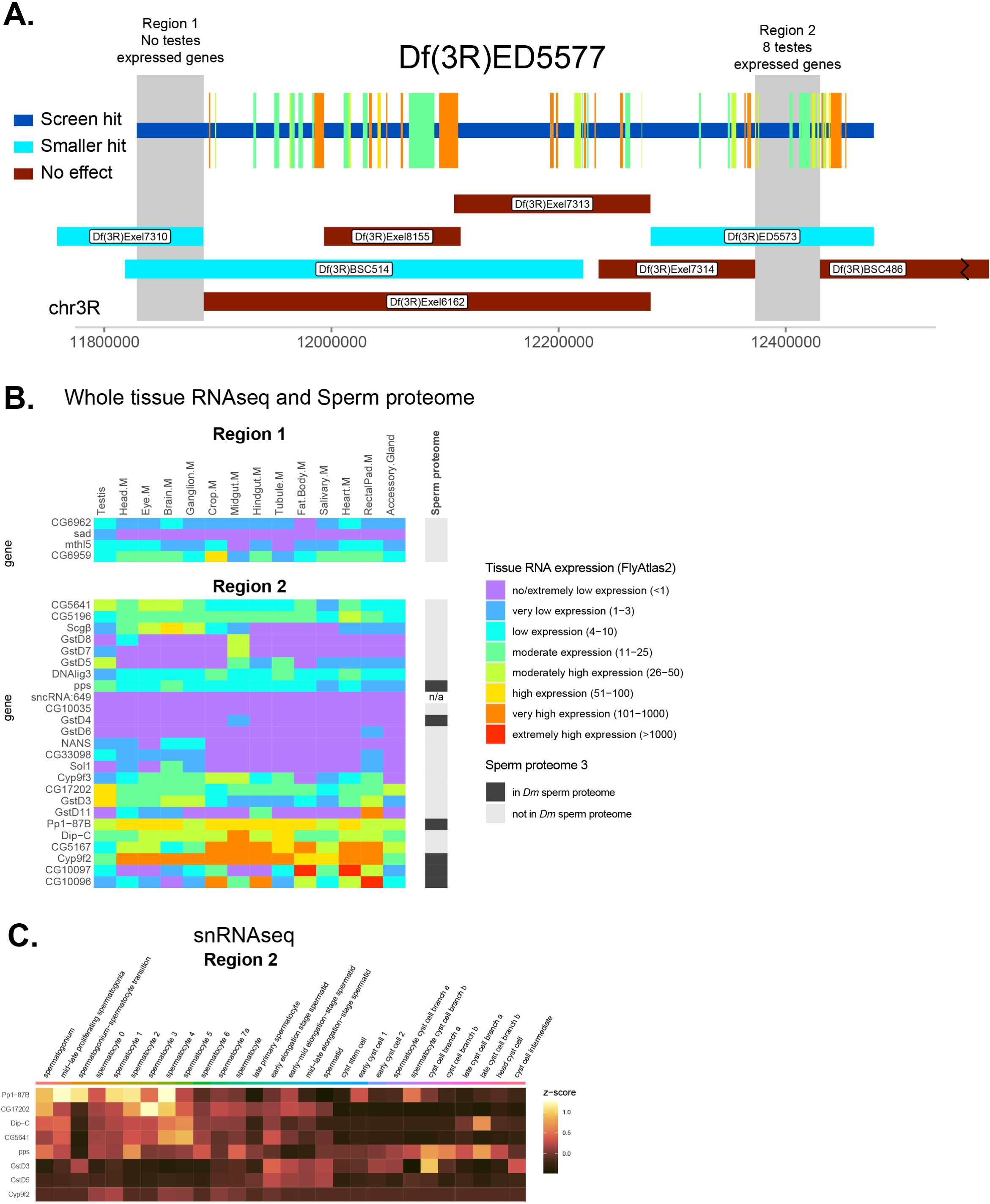
Analysis of screen hit Df(3R)ED5577. **A.** genomic region of DF(3R)ED5577. Horizontal bars represent regions of the genome removed by Df screen hits and each smaller Df tested. Interaction status color code in insets. Genes removed by the Df from the screen are represented by vertical bars with their width indicating the region of the genome they occupy. Gene color indicates expression level (FPKM) in testis from FlyAtlas 2 (legend in B). Axis is FlyBase Gene Coordinates. Grey regions represent the region of the genome discussed in the Results/Discussion. **C.** Heatmap of the FPKM from RNAseq (FlyAtlas 2) of genes in grey region in A. for various tissues. Rightmost column indicates presence or absence of proteins encoded by these genes in the sperm proteome 3. **C.** Heatmap of snRNAseq data from FlyCellAtlas of gens in C. with moderate or higher expression in the testis.

**Figure S5.**
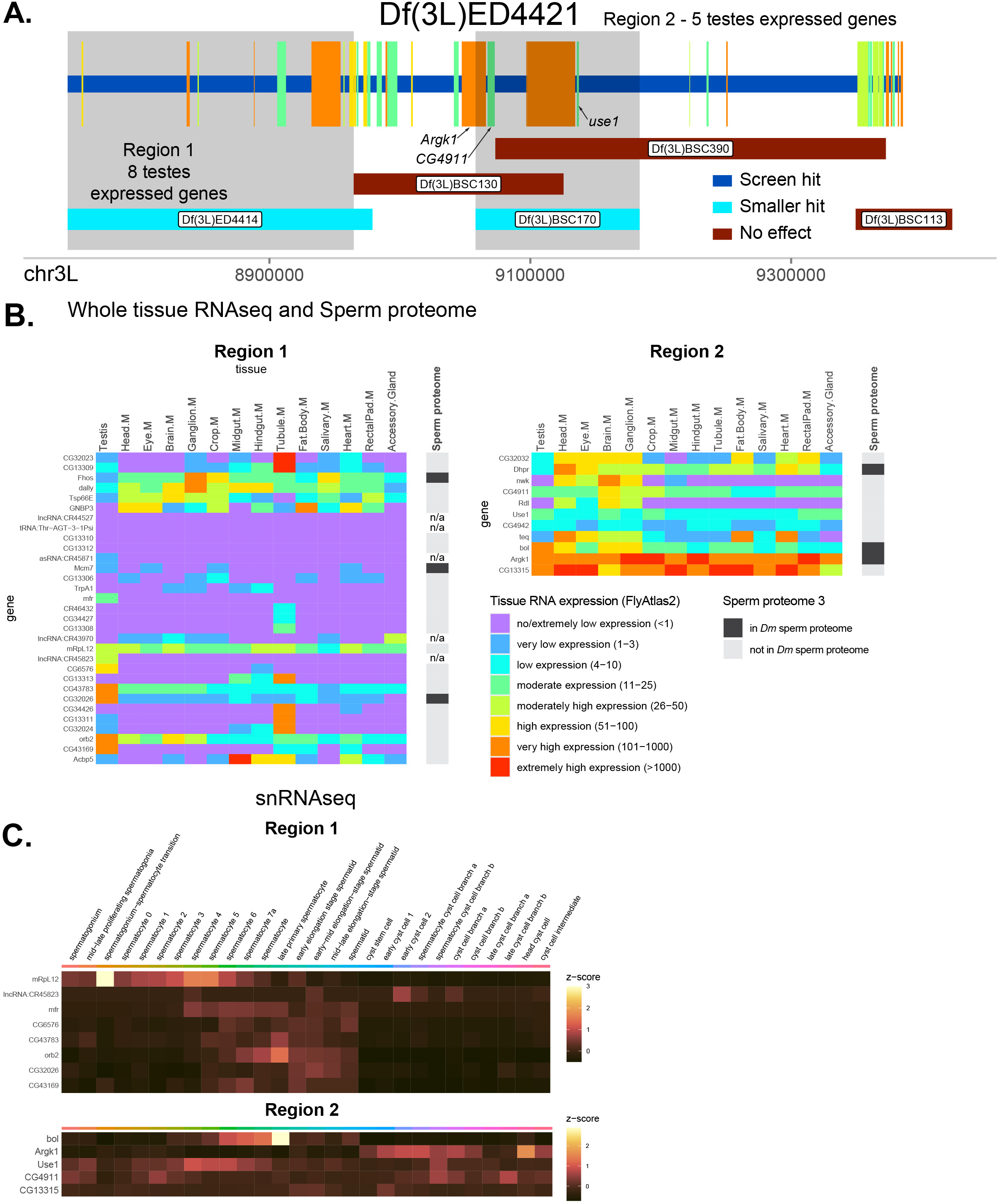
Analysis of screen hit Df(3R)ED4421. **A.** genomic region of DF(3R)ED4421. Horizontal bars represent regions of the genome removed by Df screen hits and each smaller Df tested. Interaction status color code in insets. Genes removed by the Df from the screen are represented by vertical bars with their width indicating the region of the genome they occupy. Gene color indicates expression level (FPKM) in testis from FlyAtlas 2 (legend in B). Axis is FlyBase Gene Coordinates. Grey regions represent the region of the genome discussed in the Results/Discussion and the specific genes discussed are indicated. **B.** Heatmap of the FPKM from RNAseq (FlyAtlas 2) of genes in grey region in A. for various tissues. Rightmost column contains results of presence or absence of proteins encoded by these genes in the sperm proteome 3. **C.** Heatmap of snRNAseq data from FlyCellAtlas of gens in C. with moderate or higher expression in the testis.

### Supplemental Files

#### File 1

Sheet 1. List of Dfs tested in this screen. Numbers in the first column correspond to the X-axes of the plots in Figure 2.

Sheet 2. Results of primary and secondary fertility screens. Sheet 3. Statistical information for Figure 3.

Sheet 4. Statistical information for Figure 4. Sheet 5. Statistical information for Figure 5B. Sheet 6. Statistical information for Figure S1.

#### File 2

Computational methods used in this study.

## Methods

### Fly stocks and husbandry

Control flies used in this study were *yw*, except for a few of the fertility assays where *w^1118^* or *oreR* was used. The PLP^OE^ transgenic fly, ubi-PLP::GFP, was previously described (Galletta et al., 2014). The Ana1::tdtomato strain was a generous gift of Tomer Avidor-Reiss (University of Toledo, Toledo, OH). The screen utilized the Bloomington Deficiency Kit (Cook et al., 2012; Roote and Russell, 2012); Bloomington *Drosophila* Stock Center, Bloomington, IN). All additional Dfs were obtained from the Bloomington *Drosophila* Stock Center. All flies were maintained at 25°C on standard cornmeal media.

### Fertility tests

One to three day old males were used for fertility testing. These males were individually crossed to 2, 1 - 3 day old virgin control females in individual vials. These flies were kept in the vial for 4 days before being cleared. All progeny that eclosed prior to the date on which one could expect the earliest 2nd generation flies were counted (17 days after being set).

### Primary screen

Virgin ubi-PLP::GFP/CyO females were crossed to males from strains from the Bloomington deficiency kit. F1 males from this cross were then collected for fertility tests. For 292 of the 311 strains, at least 3 individual males were tested. For the remaining 19, only 2 were tested. Due to the large number flies to be tested and an interruption caused by the COVID-19 pandemic, fertility tests were performed in several batches. A reduction in fertility in the primary screen was defined as a reduction in the average fertility of test males to less than 65% the fertility of the average fertility of control males. This is approximately 1.5 standard deviations below the mean for most batches.

### Secondary screen

Df lines that showed reduced fertility in the primary screen were retested. The secondary screen used many more individual males than the primary. Hits were determined by t test relative to control fly fertility.

### Sample preparation - Head-tail connection assay

Testes were dissected from adult males in *Drosophila* S2 media (Gibco, ThermoFisher Scientific, Waltham, MA). An individual testis was then moved to a drop of S2 media on a clean slide. Forceps were then used to grab the tip of the testis, near the stem cell niche. An insect pin was then used to open the opposite end of the testis, near where the seminal vesicle is attached. A fine glass needle was then used to squeeze the contents of the testis onto the slide starting at the closed end. Glass needles were then used to separate freed cysts from each other. A No. 1.5 coverslip was then placed on the drop and media was removed using a Kimwipe (Kimberly-Clark, Irving, TX) until cysts were slightly squashed. Slides were then treated as described (Varadarajan et al., 2016) with modification. As soon as cyst were sufficiently squashed, slides were flash frozen in liquid nitrogen, the coverslip removed using a razor blade and the slides were stored in 100% ethanol at -20°C. Samples were fixed in 4% formaldehyde in Phosphate Buffered Saline (PBS) for 20 min. Samples were blocked for at least 30 min in 5% normal goat serum (MilliporeSigma, Burlington, MA) in PBST (PBS + 0.1% Tween 20). Primary antibody (guinea pig anti-Asl; 1:10,000 (Klebba et al., 2013)) in block was incubated at room temperature for 1-2 hours. Samples were washed with 3 changes of PBST. Secondary antibody (anti-guinea pig Alexa Fluour 647; 1:1000; ThermoFisher Scientific) with DAPI (1:1000; ThermoFisher Scientific) in block was incubated for 1 hour at room temperature. Slides were then dunked 10 times in PBS. A drop of Aquapolymount (Polysciences, Warrington, PA) was placed over the sample and cover with a No. 1.5 coverslip.

### Microscopy

Slides were imaged on a Nikon Eclipse Ti2 microsope (Nikon Instruments, Melville, NY) with - a W1 spinning disk confocal head (Yokogawa, Life Science, Tokyo, Japan); 405-, 488-, 561-, and 641-nm laser lines; and Prime BSI cMOS camera (Teledyne Photometrics, Tucson, AZ); a 100X/1.49 TIRF oil immersion objective. Images were collected using Nikon Elements software (Nikon Instruments). Images were analyzed and processed using FIJI (Image J; National Institutes of Health, Bethesda, MD).

### Quantification and statistical analysis Head Tail-Connection Assay

Leaf and early canoe stage cysts were selected for analysis. Cysts were only analyzed if it appeared that the entire cysts, and therefore all or most of its nuclei, was able to be analyzed. Nuclei were scored as attached if the centriole adjunct, as marked by Asl, was is close apposition to the DAPI stained nucleus. This scoring was used even if the point of attachment of the centriole to the nucleus appeared abnormal. Thus, this is a conservative measurement that likely underestimates the extent of HTCA defects. 10 testes of each genotype were examined. The fraction of nuclei without an attached centriole is presented.

### Docking angle and centriole/nuclear distance

Analysis of docking angle and centriole/nuclear distance was done on round spermatids (RSTs) as described (Galletta et al., 2020). Centrioles with their long axis almost parallel to the imaging plane were selected and Z-projections to include the centriole and the nucleus were generated. The perimeter of the nucleus, as defined by DAPI staining, was selected by hand in ImageJ and the centroid was determined. The point of the centriole closest to the nucleus and farthest from the nucleus were selected. The distance between the nuclear centroid and the closest end of the centriole was the “centriole/nuclear” distance. The angle between these three points (centroid, near centriole, far centriole) was then calculated and was the “docking angle”. As previously noted, both of these measurements likely underestimate the severity of defects because they measure the 2D arrangement of objects arranged in 3D.

### Statistical Analysis

All comparisons were made with unpaired t test, with Welch’s correction when appropriate. Sample sizes are presented in supplementary tables or in figure legends. In all cases the mean ± standard deviation is presented. Data analysis was performed with Execl (Microsoft, Redmond, WA) and Prism (Graphpad, Boston, MA).

### Computational analysis of gene expression datasets

Expression levels and patterns of genes within the selected genomic regions of interest were investigated on two levels. First, we utilized RNA-sequencing data from FlyAtlas2 (Leader et al., 2018) to examine adult tissue-specific gene expression levels. For this analysis, we used the most recent gene count table (Gillen, A. (2023.11.14) FlyAtlas2 2023 Data) and adopted the FPKM thresholds employed in FlyBase to categorize gene expression levels. Next, we filtered for genes with “moderate” or higher expression levels (≥11 FPKM) in adult testes (hereafter referred to as “testes-expressed genes”) and investigated their expression across individual testes cell types using single-nucleus RNA-sequencing data from Fly Cell Atlas (Li et al., 2022) and the Seurat toolkit for analysis (Hao et al., 2024). For the single-nucleus RNA-seq analysis, we used the integrated 10x and SMART-seq2 testis datasets excluding fat body, hemocyte, muscle, epithelium, tracheal, pigment cell and “unknown” cell types. To illustrate expression patterns of selected genes, their average scaled expression (z-score) was calculated for each cell type. Identification of genes present in the sperm proteome (Garlovsky et al., 2022) was done manually.

We plotted the number of testes-expressed genes removed each deletion (box plot) against the deletion’s phenotype (‘screen hit’ vs ‘screen miss’). For a more detailed representation, we visualized these genes as color-coded segments over selected regions, colors representing expression levels in the testes based on FlyAtlas2 data. Gene coordinates were obtained from the FlyBase annotation. Detailed computational methods and code are available in the supplemental code file accompanying this paper.

